# Differential genetic strategies of *Burkholderia vietnamiensis* and *Paraburkholderia kururiensis* for root colonization of *Oryza sativa* ssp. *japonica* and ssp. *indica*, as revealed by Tn-seq

**DOI:** 10.1101/2022.04.14.488431

**Authors:** Adrian Wallner, Nicolas Busset, Joy Lachat, Ludivine Guigard, Eoghan King, Isabelle Rimbault, Peter Mergaert, Gilles Béna, Lionel Moulin

## Abstract

*Burkholderia vietnamiensis* LMG10929 (*Bv*) and *Paraburkholderia kururiensis* M130 (*Pk*) are bacterial rice growth-promoting models. Besides this common ecological niche, species of the *Burkholderia* genus are also found as opportunistic human pathogens while *Paraburkholderia* are mostly environmental and plant-associated species. Here, we compared the genetic strategies used by *Bv* and *Pk* to colonize two subspecies of their common host, *Oryza sativa ssp. japonica* (cv. Nipponbare) and *ssp. indica* (cv. IR64). We used high-throughput screening of transposon insertional mutant libraries (Tn-seq) to infer which genetic elements have the highest fitness contribution during root surface colonization at 7 days post inoculation. Overall, we detected twice more genes in *Bv* involved in rice roots colonization compared to *Pk*, including genes contributing to the tolerance of plant defenses, which suggests a stronger adverse reaction of rice towards *Bv* compared to *Pk*. For both strains, the bacterial fitness depends on a higher number of genes when colonizing *indica* rice compared to *japonica*. These divergences in host pressure on bacterial adaptation could be partly linked to the cultivar’s differences in nitrogen assimilation. We detected several functions commonly enhancing root colonization in both bacterial strains e.g., Entner-Doudoroff (ED) glycolysis. Less frequently and more strain-specifically, we detected functions limiting root colonization such as biofilm production in *Bv* and quorum sensing in *Pk.* The involvement of genes identified through the Tn-seq procedure as contributing to root colonization i.e., ED pathway, c-di-GMP cycling and cobalamin synthesis, was validated by directed mutagenesis and competition with WT strains in rice root colonization assays.

**Importance:** Burkholderiaceae are frequent and abundant colonizers of the rice rhizosphere and interesting candidates to investigate for growth promotion. Species of *Paraburkholderia* have repeatedly been described to stimulate plant growth. However, the closely related *Burkholderia* genus hosts both beneficial and phytopathogenic species, as well as species able to colonize animal hosts and cause disease in humans. We need to understand to what extent the bacterial strategies used for the different biotic interactions differ depending on the host and if strains with agricultural potential could also pose a threat towards other plant hosts or humans. To start answering these questions, we used here transposon sequencing to identify genetic traits in *Burkholderia vietnamiensis* and *Paraburkholderia kururiensis* that contribute to the colonization of two different rice varieties. Our results revealed large differences in the fitness gene sets between the two strains and between the host plants, suggesting a strong specificity in each bacterium-plant interaction.

## Introduction

Species of the *Burkholderia* and closely related *Paraburkholderia* genera are highly prolific rhizosphere colonizers (1, 2). Their persistence and competitiveness in the rhizosphere environment can be explained by strong secondary metabolite production as well as efficient nitrogen cycling, mineral solubilization and phytohormone biosynthesis (3–5). Furthermore, several *Paraburkholderia* and at least one *Burkholderia* species can fix atmospheric nitrogen. Two well studied models, *Paraburkholderia kururiensis* strain M130 (hereafter called *Pk*) and *Burkholderia vietnamiensis* strain LMG10929 (hereafter called *Bv*) demonstrate strong rice root colonization, endophytic lifestyles and significant plant growth promotion through transfer of fixed nitrogen (6–9). Despite this convergence in their plant beneficial features, both bacteria belong to distinct genetic backgrounds. While *Paraburkholderia* species are often found in beneficial relationships and symbioses with plants (10–12), *Burkholderia* members comprise human pathogens and opportunists as well as fungal and plant pathogens (13).

Bacterial genes used for plant colonization have been screened in several model bacteria but also on a broader scale using microbiome approaches to reveal plant-associated functions (1). A few studies have compared plant-adapted *Burkholderia s.l. (sensu largo,* e.g. the former genus that is now regrouping the newly defined *Burkholderia*, *Paraburkholderia*, *Caballeronia* and others) at the genomic level (4, 11), or profiled the transcriptome of bacteria stimulated by root exudates (14, 15). However, there is no record of a comparison between the strategies used by plant-adapted *Burkholderia* and *Paraburkholderia* species. The host plant genotype’s impact on bacterial colonization strategies also remains poorly explored in these bacterial genera. Rice is an interesting model to assess *Burkholderia s.l.* adaptation to the plant environment as it is hosting plant growth promoting model strains from both *Burkholderia* and *Paraburkholderia* genera. It was repeatedly demonstrated that rice genotypes influence the composition of their microbiome at the rhizosphere and rhizoplane levels (16, 17). In particular, a study on 95 different *Oryza sativa* subsp. *indica* and subsp. *japonica* varieties showed significant differences in microbiome composition between both rice subspecies, which was related to their nitrogen use efficiency and the presence of a particular nitrate transporter in indica varieties (18).

Transposon mutagenesis sequencing (Tn-seq), is a high-throughput screening method that combines transposon insertional mutagenesis followed by sequencing of the insertion sites (19). It is a powerful tool that leads to immediate identification of genes of interest improving or reducing the bacteria’s fitness in a tested condition. This methodology has been successfully used to unravel important genes and functions in plant-pathogenic or plant-symbiotic interactions (20–22). Commonly between pathogens and mutualists, genes functioning in amino acid and purine metabolism as well as in cell motility were detected to be required for root colonization (21, 23, 24).

In the present study, we used Tn-seq, to perform a genome-wide identification of genes involved in rice-root colonization in *Bv* and *Pk*. In detecting which genes influence the fitness of *Bv* and *Pk* we aim at unraveling their commonalities and differences in root colonization strategies. We also analyzed the association strategies of *Bv* and *Pk* with the two rice genotypes *Oryza sativa* subsp. *japonica* (cv. Nipponbare) and *indica* (cv. IR64) to understand how the host-factor can influence colonization strategies. Overall, we identified a total of 1,404 and 540 genes that influence the fitness of *Bv* and *Pk* respectively during rice root colonization. Our results underline the importance of motility, amino acid and vitamin metabolism, stress response as well as biofilms for the efficient association of these bacteria with rice roots.

## Results

### Quality of Tn-Seq libraries and essential genomes in *Pk* and *Bv*

To generate a genome-wide library of insertion mutants for *Pk* and *Bv* strains we used a mariner transposon that targets genomic thymine-adenosine (TA) sites (**Materials and methods**). The genome of *Bv* contains 95,075 TA sites and we estimated the mutant population at 1.6 x 10^8^ cfu, which represents a 1683x coverage of the total TA sites. The saturation level of the *Pk* library was lower although still significant at a 38x coverage given the 4.0 x 10^6^ cfu obtained after transposon mutagenesis for a total of 106,136 total TA sites contained in the genome.

To further assess the quality of the Tn-seq libraries we determined the essential genomes required by both bacteria for optimal development in a rich liquid growth medium. Both bacteria have similar genome sizes (6,820 and 6,436 genes for *Bv* and *Pk* respectively) and comparable proportions of genes (661 or 9.7% and 638 or 9.9%, respectively) that are required for optimal growth in a rich, liquid medium (**Figure 1A; Table S1**). The size of the essential genomes determined in a controlled liquid medium are in the order of magnitude of what has been observed for other prokaryotes, including *Burkholderia spp.* (26–30).We used the Minimal Gene Set tool implemented in the MicroScope platform (**Materials and methods**) to extract a core list of the predicted minimal bacterial gene sets from *Bv* and *Pk*. On a total of 206 core bacterial genes defined by their conservation among multiple bacterial genomes (25), 151 and 150 were identified as essential by our approach in *Bv* and *Pk,* respectively. In 10 and 9 cases of genes belonging to the minimal gene set but predicted as non-essential by our approach in *Bv* and *Pk* respectively, there are duplicates present in the genome.

**Figure 1.**
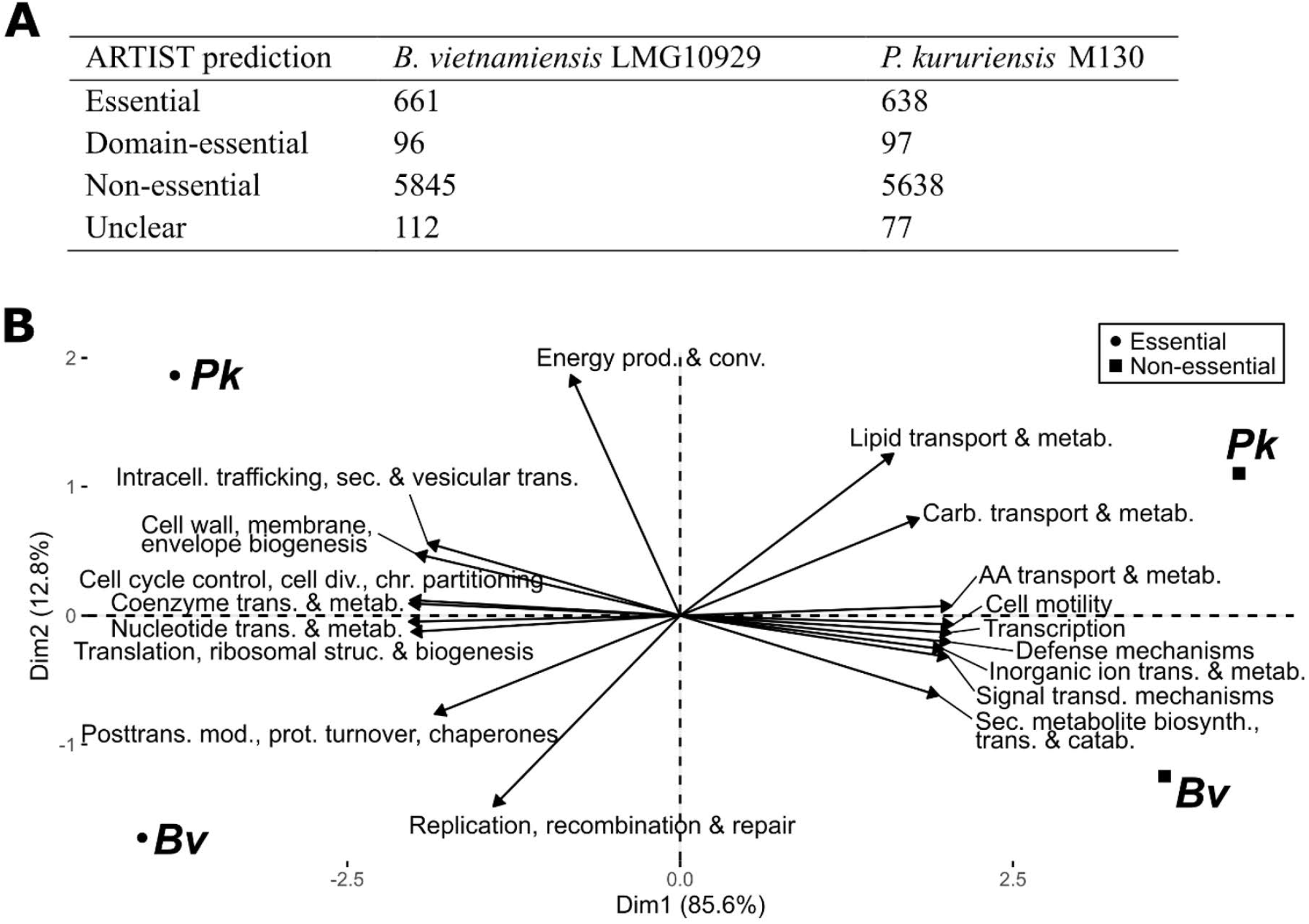
Distribution of genes significantly impacting bacterial fitness in a rich liquid medium. (A) Number of genes predicted to be essential, domain-essential (only insertions in part of the gene lead to abundance decrease) or non-essential by the EL-ARTIST pipeline. (B) Principal component analysis of essential and non-essential genes of Bv and Pk in a rich liquid medium. The genes were classified according to their COG categories and the PCA was built in regard to the abundance of these categories.

According to their distribution in COG categories, little variance differentiates the essential genomes of *Pk* and *Bv* (**Figure 1B**). As expected in a liquid, rich medium, genes involved in motility, defense and nutrition are largely unessential. Conversely, structural components of the cell and the general replicative machinery are predictably essential (**Table S1**).

Given the strong saturation level of the TA insertion sites and the coherent essentiality results observed in the control setting, we can safely assume that both mutant libraries allow a reliable analysis of the impact of genes on the bacterial fitness.

### Colonization of the two rice genotypes by *Pk* and *Bv*

Prior to genetic analyses, we assessed the colonization efficiencies of *Bv* and *Pk* on Nipponbare *(japonica)* compared to IR64 *(indica)* rice genotypes (**Figure 2**). Overall, both bacteria display a similar colonization dynamic with an increasing population density during the first week and a decrease in the second week. A host-genotype effect is observed in the colonization phenotype displayed by *Pk* as significantly different root colonizing populations are observed on IR64 and Nipponbare at 3 dpi and 14 dpi (**Figure 2**). *Bv* on the other hand displays a steady colonization pattern between both plant genotypes at all measured time points. These observations confirm that the rice-colonizing populations of *Bv* and *Pk* can be compared, especially at 7 dpi, which was selected for further analyses. In our following analyses, we consider that the bacterial adaptations observed result purely from root surface colonization, as the endophytic populations at 7 dpi are inferior by several log scales to the surface colonizing population (6).

**Figure 2.**
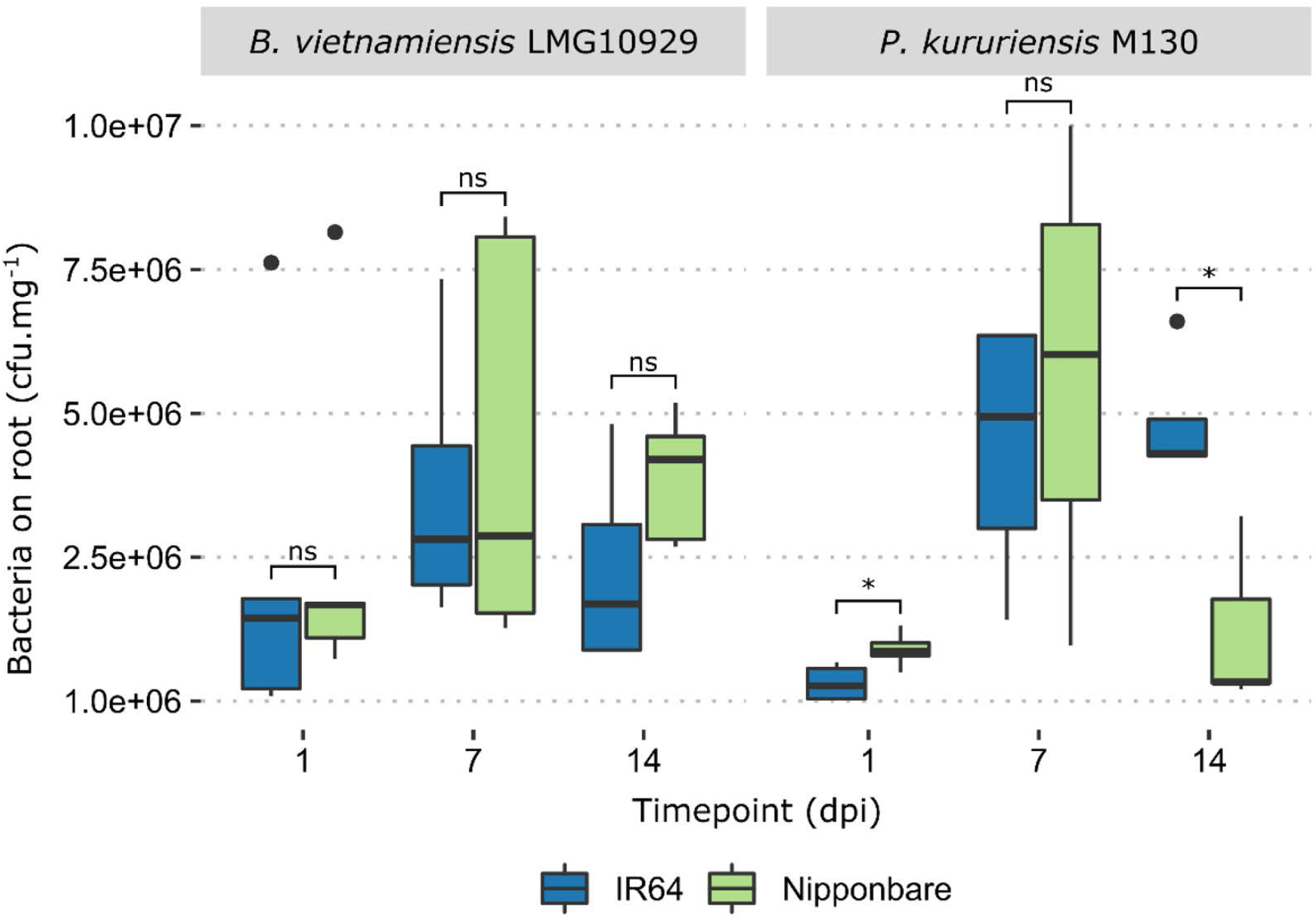
Rice colonization efficiencies for *Bv* and *Pk.* Nipponbare and IR64 rice seedlings were inoculated at 5 days post germination with 1.10^7^ bacterial cells, total root systems were harvested at different time points and the associated bacterial population was estimated (p<0.05, Wilcoxon test). The boxplots were generated using 5 replicates for each condition. Outliers are represented as black dots.

### Identification of Rice colonization genes

To assess gene fitness for rice root colonization, and its host-dependent variation in the two model bacteria, we inoculated the Tn-Seq mutant libraries on Nipponbare and IR64 rice genotypes. Seven days after inoculation, we harvested and pooled five root systems per replicate, representing a total of 1.2 x 10^7^ colonization events (**Figure 2**) and a more than 100-fold coverage of the available mutant diversity. We performed a first Tn-seq analysis by pooling the reads of IR64 and Nipponbare isolates together to infer the genes globally required for the association with rice. The read frequencies were compared to the control condition to establish a root fitness score for each gene. In this manner, we identified 1,404 and 540 genes as significantly impacted (enriched or depleted) after root-colonization by *Bv* and *Pk*, respectively (**Figure 3A**). Colonization-depleted genes will be our major focus as they are positively associated with bacterial fitness. Inversely, colonization-enriched genes diminish the bacterial fitness during root colonization. We organized the identified colonization genes according to their general function based on their clusters of orthologous groups (COG) annotation (**Figure 3B**). Colonization-enriched and -depleted genes share a similar distribution with amino acid metabolism, cell wall/membrane/envelope biogenesis, transcription and cell motility being amongst the most frequent categories, consistent with the expected implication of nutrition, motility and morphological adaptations involved in plant colonization.

**Figure 3.**
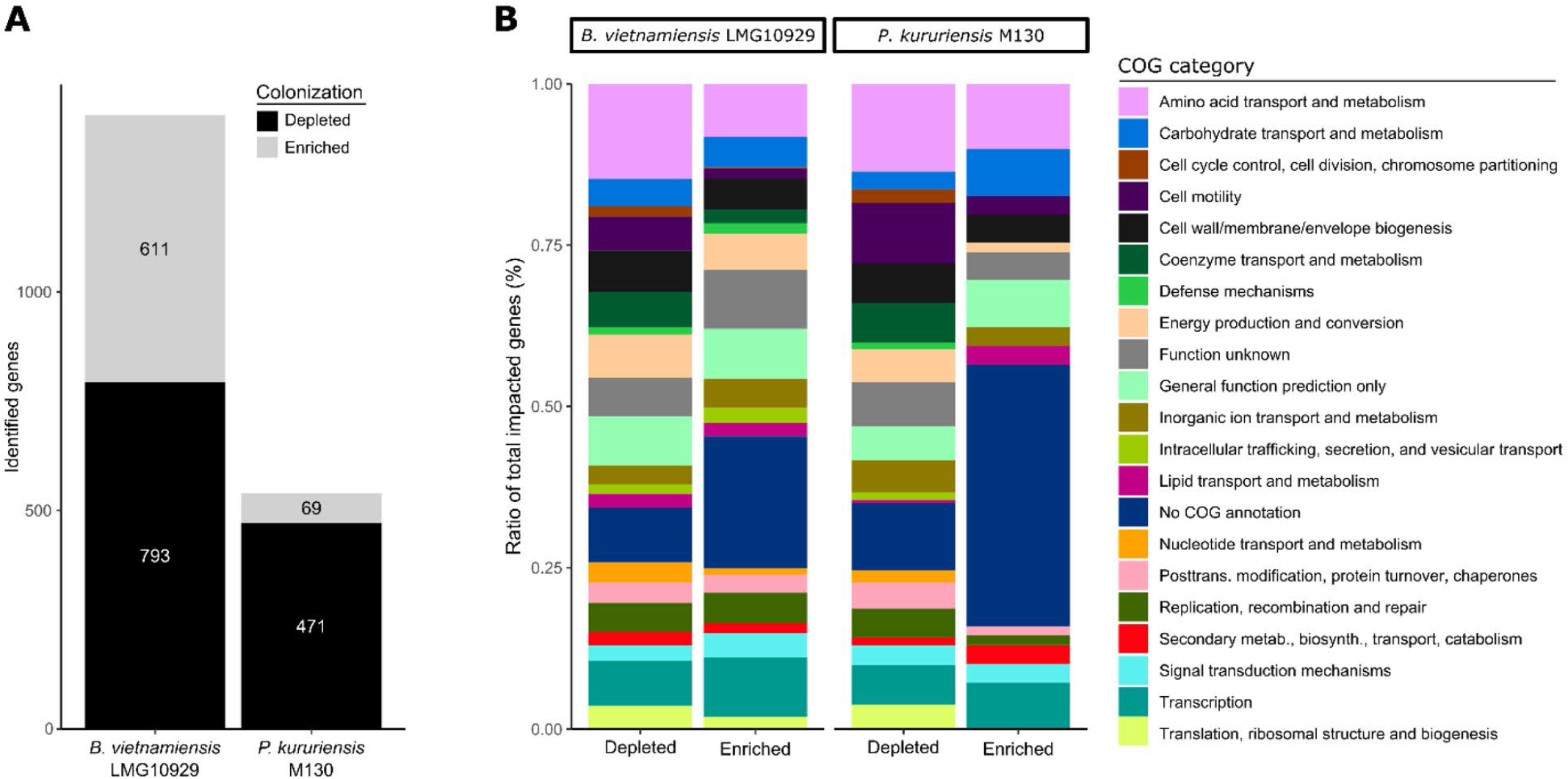
Distribution of genes significantly impacting bacterial fitness during rice colonization. Colonization events on IR64 and Nipponbare rice cultivars were pooled for this analysis. (A) Transposon insertions in genes leading to a 1.5-fold or higher decrease in abundance on roots at padj < 0.05 were identified to be colonization-depleted. Those leading to a 1.5-fold or higher abundance were identified to be colonization-enriched. (B) Distribution of genes significantly depleted or enriched during rice colonization along COG categories in ratio of total impacted genes.

We identified a total of 2,071 core gene families for *Bv* and *Pk*, sharing homologues in both strains (**Table S2**). In *Bv* and *Pk,* 192 genes, respectively 24% and 41% of the colonization-depleted genes are part of the core-genome and equally depleted in both strains (**Figure 4**). *Bv* displays 68% more colonization-depleted genes than *Pk* (**Figure 3**). Interestingly, a majority of these *Bv* specific colonization-depleted genes (53%) are part of the core genome (**Figure 4**). Similarly, a lower, although substantial portion of the genome shared with *Bv* (37%), is required by *Pk* specifically for efficient root colonization (**Figure 4**). Thus, although a large portion of the core genes contribute to root colonization in both strains, many core genes are only required for colonization in one or the other strain, indicating that the impact of these genes on root colonization is dependent on the genome context.

**Figure 4.**
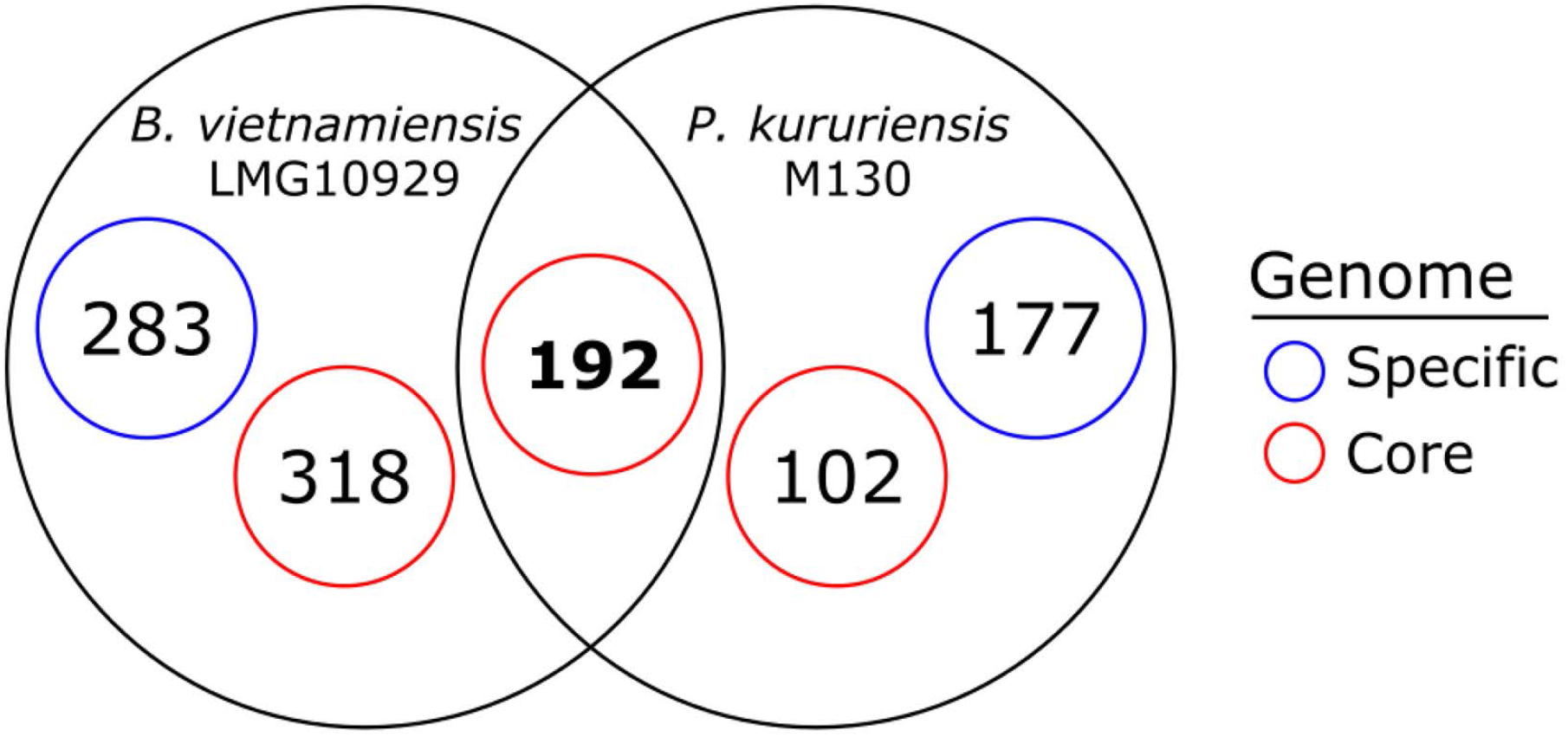
Distribution of colonization-depleted genes along the core- and specific-genomes of *Bv* and *Pk*. There are a total of 2072 genes that present significant homologies in the genomes of *Bv* and *Pk* and thus considered part of the core-genome (Materials and methods). A fraction of this core-genome is colonization-depleted in both species. Additionally, a substantial proportion of the genes that are found to be colonization depleted in a species or the other also belong to the core-genome.

### Rice cultivar dependent colonization specialization

Next, we performed a separate analysis of the reads for each rice cultivar and then compared the lists of significantly colonization-depleted and -enriched genes to infer cultivar-specificities. For both strains, the majority (~65%) of identified genes impacted the bacterial fitness on both IR64 and Nipponbare rice (**Table 1**). However, both bacteria displayed a greater number of colonization-depleted genes (41% and 49% higher for *Bv* and *Pk* respectively) on IR64 compared to Nipponbare rice (**Table 1**).

**Table 1.** Distribution of genes significantly impacting bacterial fitness on IR64 and Nipponbare rice. Colonization events on IR64 and Nipponbare rice cultivars were analyzed separately.

| Strain | Colonization impact | Common | IR64 | Nipponbare | Total |
| --- | --- | --- | --- | --- | --- |
| *B. vietnamiensis*<br>LMG10929 | Enriched | 507 | 83 | 130 | 720 |
|  | Depleted | 635 | 225 | 134 | 994 |
|  | **Total** | **1142** | **308** | **264** | **1714** |
| *P. kururiensis*<br>M130 | Enriched | 48 | 57 | 36 | 141 |
|  | Depleted | 374 | 98 | 50 | 522 |
|  | **Total** | **422** | **155** | **86** | **663** |

Core genes of *Bv* and *Pk* that are colonization-depleted on both cultivars belong primarily to housekeeping categories such as central metabolism, cell cycle control and motility (**Figure 5A)**. The additional genes, depleted during IR64 colonization are chiefly attributed to amino acid metabolism, transcription, cell wall/membrane/envelope biogenesis and energy production in both bacterial strains (**Figure 5B & 5C**). Interestingly, contrary to the global trend, in *Pk* the COG categories “ replication, recombination & repair”, “coenzyme transport & metabolism”, “carbohydrate metabolism”, and “inorganic ion transport” are more strongly impacted on Nipponbare than IR64 (**Figure 5C**). The same trend is observed in *Bv* for the COG categories “signal transduction mechanisms” and “coenzyme transport and metabolism” (**Figure 5B**).

**Figure 5.**
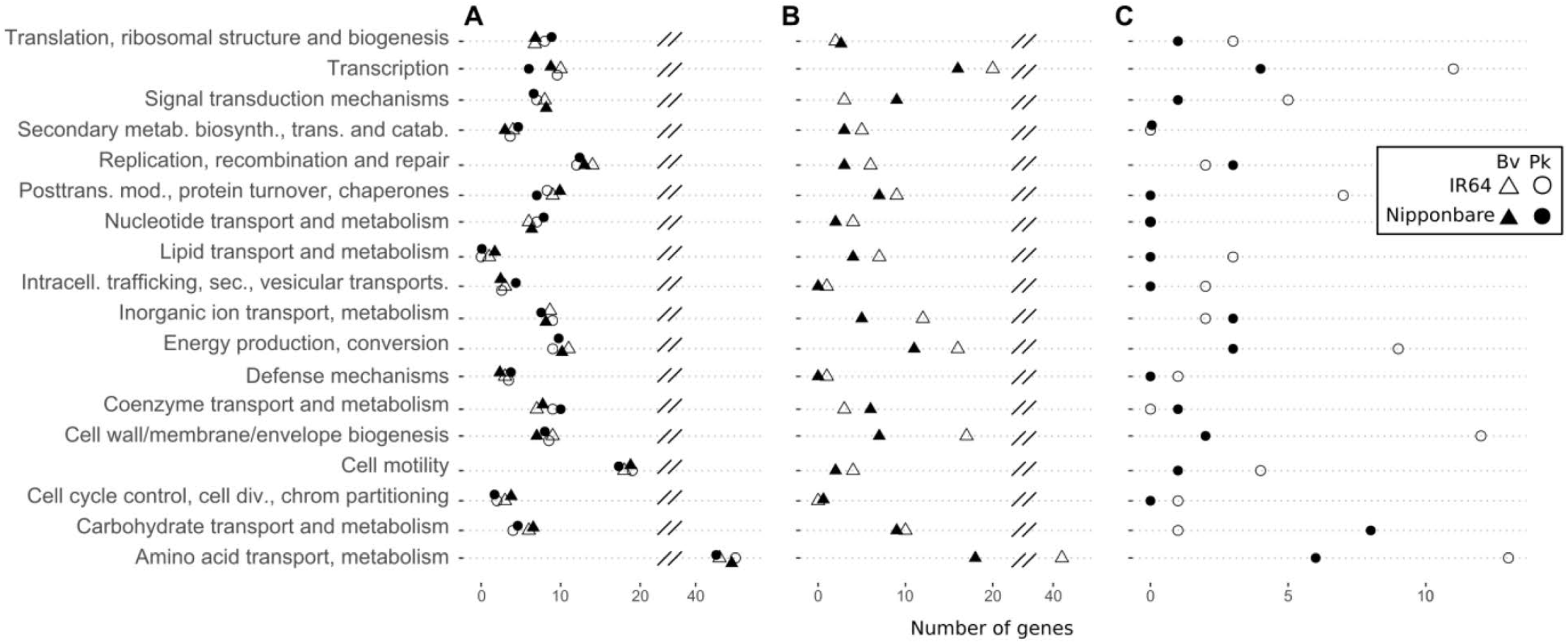
Distribution of colonization-depleted genes on IR64 and Nipponbare rice cultivars. Distribution along COG categories of: (A) core genes of *Bv* and *Pk* that are colonization-depleted on both rice cultivars, (B) genes that are specifically colonization-depleted in *Bv* and (C) genes that are specifically colonization-depleted in *Pk*.

### Tn-seq reveals bacterial functions involved in early rice colonization

Many genes significantly impacting the colonization fitness of *Bv* and *Pk* are clustered together within operons supporting the validity of our results. In these cases, there is furthermore a strong conservation of either gene enrichment or depletion within the same operon. In several cases explored hereafter, the lack of detection of a complete operon can be explained by the presence of homologues for some of the genes resulting in functional complementation.

### Functions required for bacterial fitness during rice-root colonization

As expected in this kind of colonization assay, mutants affected in motility and chemotaxis functions (*flg, flh, fli, mot)* were depleted from the root colonizing population (**Table 2 & Table S3**). For both bacteria, we further detected many genes involved in amino acids *(arg, his, ilv, leu, lys, met, pro, ser* and *trp*) and nucleotide synthesis *(pur and pyr)* that, when mutated, negatively impacted the bacteria’s fitness on plants. Genes involved in the synthesis of enzymatic cofactors for amino acid metabolism such as vitamin B1 (thiamin) were also colonization-depleted in both strains. Multiple Tn-seq studies reported that auxotrophy for certain amino acids is disadvantageous for root colonization and can limit plant growth promotion and biocontrol potential (23, 31).

**Table 2.**
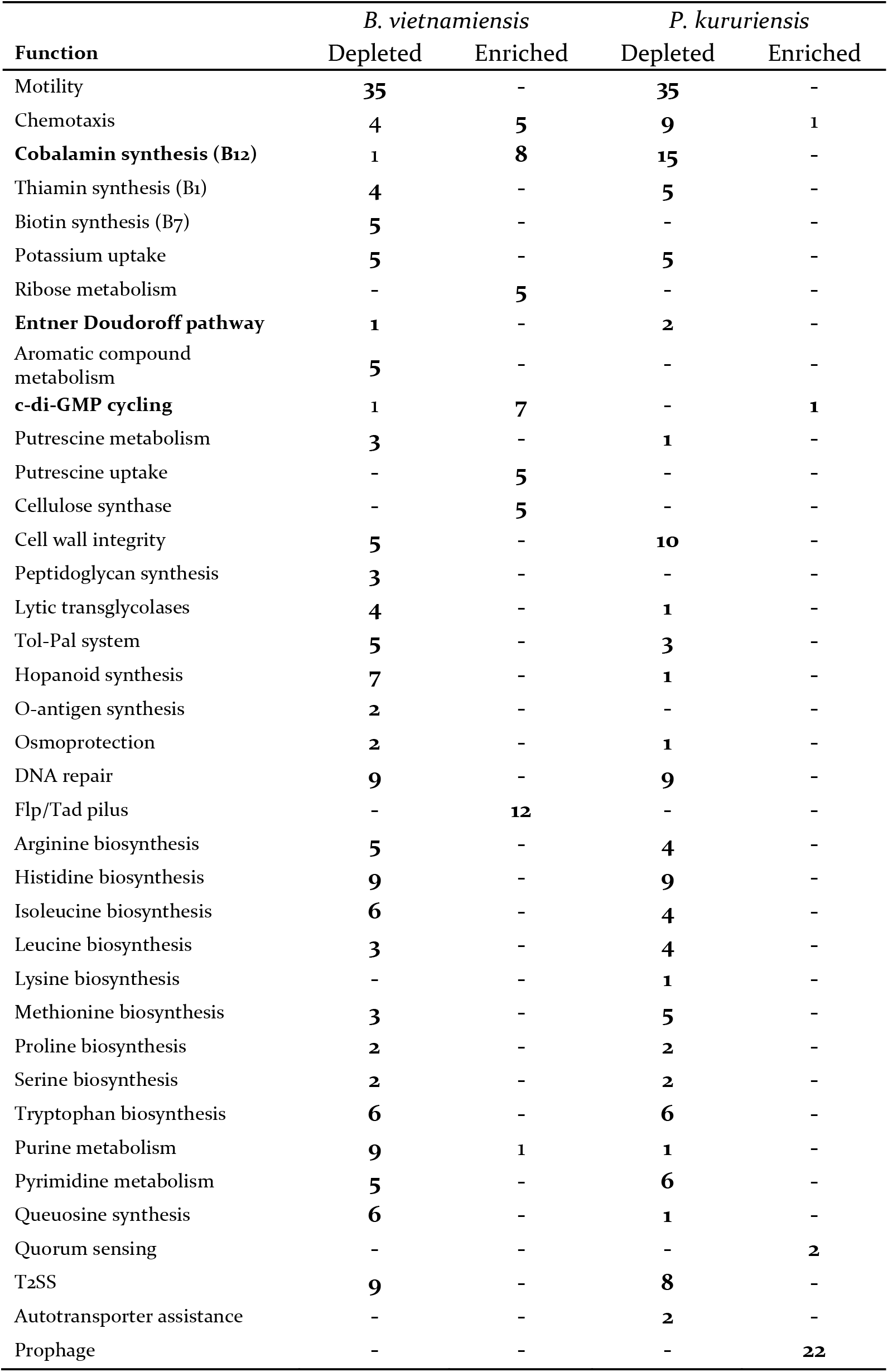
Selection of functions and number of corresponding genes impacting rice colonization efficiencies when mutated. The detailed list of genes is provided in Table S3.

Additionally, genes involved in potassium nutrition (*kdpA-F*) were similarly depleted. Mutants of both *Bv* and *Pk* affected in central elements of the Entner-Doudoroff glycolysis (ED) pathway (*edd, zwf*) suffered a significant fitness decrease on rice. This pathway is involved in the metabolism of gluconate which is not a dominant sugar of rice exudates (32) suggesting an alternative role than sole sugar assimilation for this pathway. The activation of the ED pathway was suggested to be a tolerance strategy towards oxidative stress through the generation of NADPH as essential cofactor for thioredoxins (33, 34).

Genes of the type 2 secretion system (T2SS; *gspD-M)* are the only ones belonging to a macromolecular secretory pathway to be colonization-depleted in both bacteria. While *Bv* possesses a single T2SS, *Pk* bears two copies (4) out of which only one is colonization-depleted suggesting that these systems are not redundant but are used differently in specific conditions.

Several genes involved in DNA maintenance and reparation, i.e., *ruvABC, xerCD* and *recABCD* were colonization-depleted in both *Bv* and *Pk* while their absence was tolerated in the relatively stress-free control medium (**Table S1**). Thus, rice appears to be inflicting considerable genotoxic stress during the process of colonization. Furthermore, the presence of osmotic stress is exemplified by the depletion of *Bv* and *Pk* mutants involved in the synthesis of the osmoprotectant trehalose *(otsAB).*

*Bv* seems to suffer additional stress as multiple functions maintaining cell wall and membrane integrity are colonization-depleted. We detected several genes involved in hopanoid synthesis (*hpnDEFHJKN*), peptidoglycan synthesis (*murAI*) and maintenance (*tolAQR, pal*) as well as lytic murein transglycosylation (*mltA, rlpA* and *mtgA*) to be depleted specifically in *Bv* mutant populations. This indicates that *Bv* has an increased requirement to maintain its cellular integrity compared to *Pk*. Consistent with an increased need in membrane maintenance, the loss of vitamin B7 (biotin) synthesis genes (*bioABCDF*) had a negative impact on the colonization of *Bv*. Biotin is a cofactor for many enzymes, especially those involved in fatty acid biosynthesis and amino acid metabolism (35, 36). Roots are known to secrete aromatic phenolic compounds which are toxic to various soil microbes. Accordingly, *Bv* mutants were depleted during colonization when impacted in genes of the ß-ketoadipate pathway (*pcaBCDK*) which allow to metabolize 4-hydroxybenzoate and protocatechuate. A final sign of the stress *Bv* is exposed to during colonization is found in the depletion of mutants for the queuosine synthesis pathway (*queACEF*, *tgt*). This hypermodified nucleoside improves translation accuracy, a need that only arose in *Bv* during rice colonization.

While no genes annotated as coding for autotransporter proteins (T5SS) are identified through our approach, the genes *tamAB* were colonization-depleted in *Pk*. TamA and TamB can be involved in outer membrane assembly, allowing surface structuration which is essential for adhesion and host-invasion by bacteria in several pathogenic models (37). Genes for vitamin B12 (cobalamine) synthesis are present in both *Bv* and *Pk* but were only colonization-depleted in *Pk* (*cobBDHIKLMNQSTUW*, *cbiB*, *btur*).

Intriguingly, several genes of this synthesis pathway were actually colonization-enriched in *Bv*, suggesting an adverse role in colonization for this co-factor involved in a multitude of enzymatic reactions.

### Functions detrimental for bacterial fitness during rice-root colonization

Only two genes were identified to be colonization-enriched in *Bv* and *Pk* simultaneously, one of which (AK36_1927/ANSKv1_30041) has an unknown function. The other *(pdeR)* is involved in cyclic di-GMP (c-di-GMP) cycling. In *Bv*, four other genes having a homologous role in c-di-GMP degradation (phosphodiesterases; **Table S3**) and two involved in c-di-GMP production (diguanylate cyclases; **Table S3**) were also colonization-enriched. C-di-GMP levels can play a role in colonization as they were demonstrated to influence biofilm production and motility in *Burkholderia* species (38). Interestingly, while several genes of the putrescine catabolism *(puuBD, speG)* were colonization-depleted in *Bv*, the entire cluster responsible for putrescine uptake *(potFGHI, puuP)* is colonization-enriched. Together, this suggests that putrescine accumulation has a negative impact on root colonization. An accumulation of intracellular putrescine could have an impact on biofilm production as recently suggested (23). Among additional biofilm-related colonization-enriched genes in *Bv*, we identified a cluster involved in cellulose synthesis *(bcsABCE).* Another *Bv* surface element that increases colonization efficiency when mutated is a Tad pilus *(tadABCEVZ, rcpAB, flp)* suggesting that this attachment or motility mechanism is suitable for an alternative condition than rice colonization. While many genes involved in carbohydrate metabolism in both bacteria were colonization-depleted, *Bv* mutants were colonization-enriched when impacted in ribose import and catabolism genes *(rbsABCK).*

*Pk* mutants impacted in the ability to sense the bacterial community through quorum sensing *(braR, rsaL)* were enriched during colonization as can be expected for organisms that lost the ability to sense population density and reduce growth rates accordingly. Genes belonging to a region enriched in elements of phage origin (ANSKV1_30083-120) had the same effect when mutated, indicating that the prophage might be induced by the plant colonization process and that its inactivation through mutation benefits the bacteria’s fitness.

### Rice cultivar specific adaptations

Genes which are colonization-depleted or -enriched specifically on one cultivar can be more rarely grouped in metabolic pathways and operon structures (**Table S4**). Still, single genes can have determining impacts on a strain’s ecology and metabolism. There are two genes that were colonization-depleted for both *Bv* and *Pk* on Nipponbare rice. One is an outer membrane porin of the OmpC family (AK36_1494 / ANSKv1_11218) and the other, *amtB,* is involved in ammonium uptake (**Table 3 & S4**). *Bv* mutants for genes involved in phosphate transport *(pstAC)* and glycerol *(ugpABQ)* metabolism were depleted on IR64 rice, further suggesting that the metabolic requirements of the plant force adaptations on rhizosphere microbes.

**Table 3.**
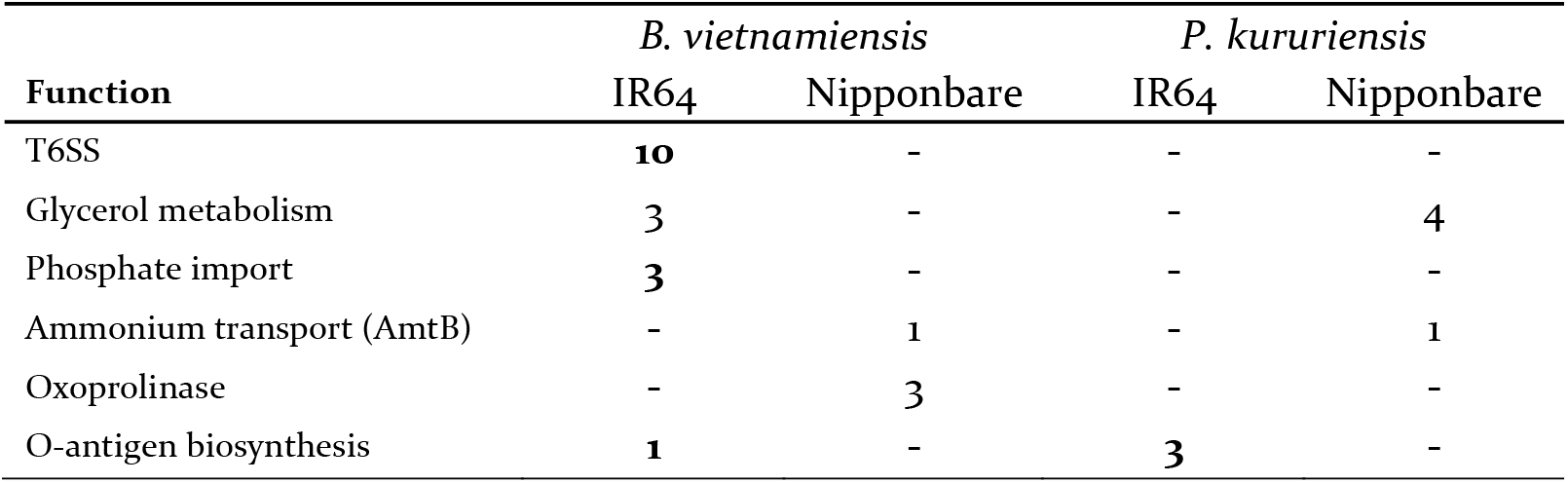
Selection of functions and number of corresponding colonization-depleted genes on IR64 and Nipponbare rice.

Several *Bv* genes involved in type 6 secretion (*tssGHIJK*) were colonization-depleted on IR64. For *Pk*, mutants impacted in O-antigen biosynthesis (*rfbABD*) were depleted on IR64. In both cases we can hypothesize that surface elements have significant and diverging repercussions depending on the type of colonized rice-host.

### Validation of three candidate genes

We underlined the importance of various genes involved in the ED pathway, c-di-GMP cycling and cobalamin synthesis among others. To validate their involvement in the colonization of rice roots, we used an insertional mutagenesis approach in *Bv* and *Pk* by targeting single copy genes that are key in the respective processes. The 6-phosphogluconate dehydratase Edd is essential for the first step of the ED pathway (39). The metal delivery protein CobW is essential for vitamin B12 synthesis (40). Finally, we chose PdeR, an enzyme with domains predicted to be involved in both c-di-GMP production and degradation.

Mutants and wild-type strains were used jointly in a competition assay and colonization efficiencies were surveyed at 7 dpi. In *Bv*, each mutation had the observable impact that had been predicted by the Tn-seq analysis (**Figure 6**). Disruption of the colonization-depleted gene *edd* reduced *Bv*’s colonization capacity while the deletion of the colonization-enriched gene *pdeR* had the opposite effect. The disruption of *cobW* had been identified in Tn-seq data as having a deleterious impact on *Pk* but did not significantly alter the root colonization capacity of *Bv*. The mutagenesis approach confirms that the disruption of *cobW* has no negative impact on *Bv* but is required by *Pk* for efficient root colonization. The only inconsistent observation occurred for the *pdeR* mutant in *Pk*, which was predicted as slightly colonization-enriched by the Tn-seq approach, but its disruption resulted in a colonization deficiency compared to the WT strain.

**Figure 6.**
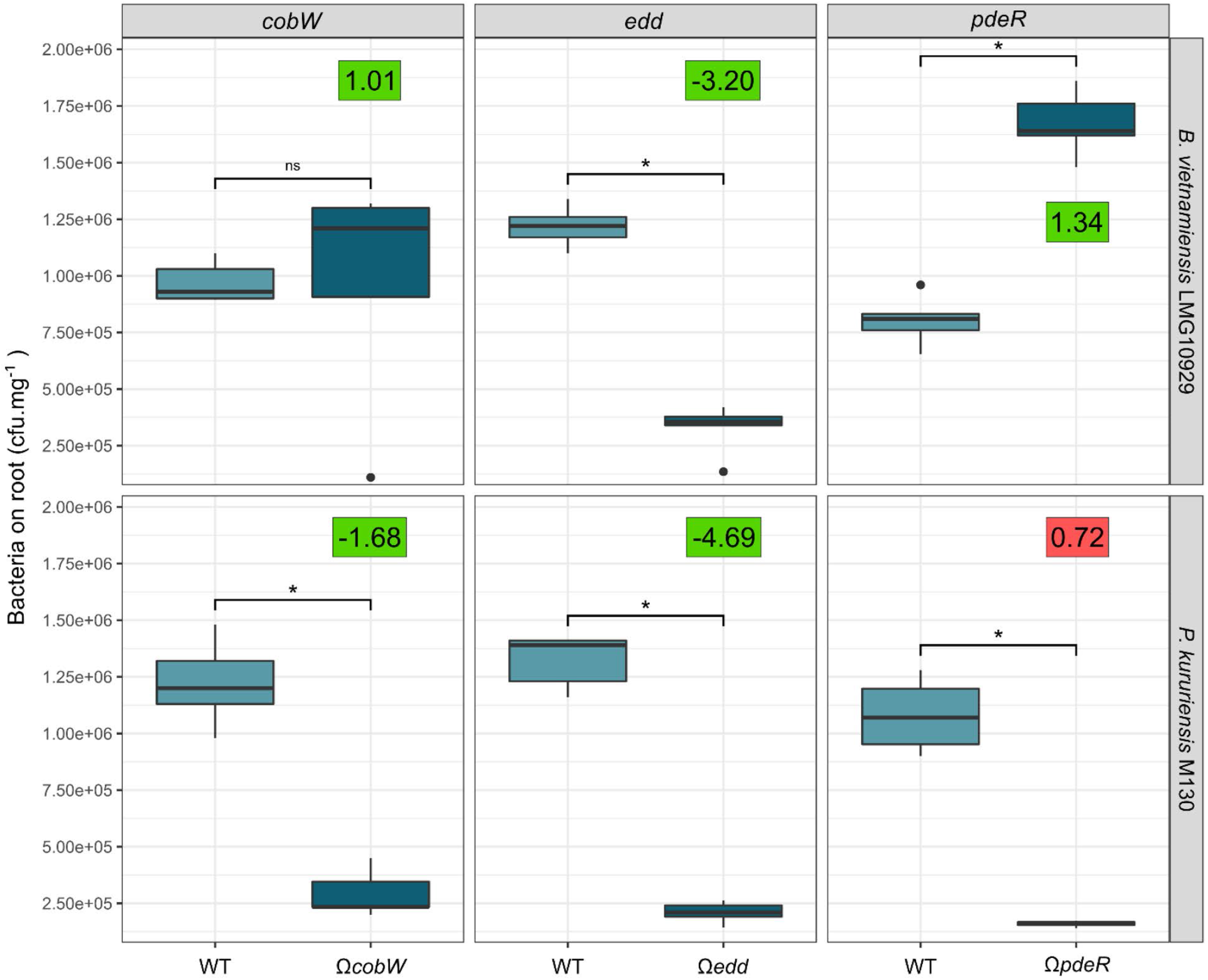
Colonization capacity of *Bv* and *Pk* insertion mutants in competition assays. Three genes, *cobW, edd* and *pdeR,* predicted by the Tn-seq approach to be involved in rice root colonization, were selected for targeted disruption in *Bv* and *Pk*. Mutants and WT strains were inoculated simultaneously on Nipponbare rice roots and enumerated at 7 dpi. Significance levels of pairwise comparisons were estimated using a Wilcoxon test (p<0.05). For each mutant, the log2 FC value observed in the Tn-seq approach on Nipponbare is displayed. Positive and negative correlations with the mutagenesis approach are expressed with green and red squares respectively.

## Discussion

Understanding which genetic bases are involved in PGPR-cereal interaction is pivotal for a controlled and informed selection of beneficial organisms and a gateway to efficient and directed strain improvement. In rice, the microbiome composition was demonstrated to be significantly modulated by the plant-genotype (16). This suggests a variance in selective pressure forced by the plant onto colonizing microbes. The present Tn-seq analysis demonstrates which genes have the strongest contribution to bacterial fitness in the early steps of rhizosphere colonization of two rice cultivars, Nipponbare and IR64 by two bacteria *Pk* and *Bv* (**Figure 7**). At this stage, we cannot exclude that additional genes might be required to colonize more mature plants as the bacteria progress and potentially reach different plant compartments. It is also known that Tn-seq approaches are insensitive to genes whose function is complemented by functional homology either within the same bacteria (gene duplicates, paralogues) or by the surrounding bacterial community, e.g., secretion systems and secreted molecules.

**Figure 7.**
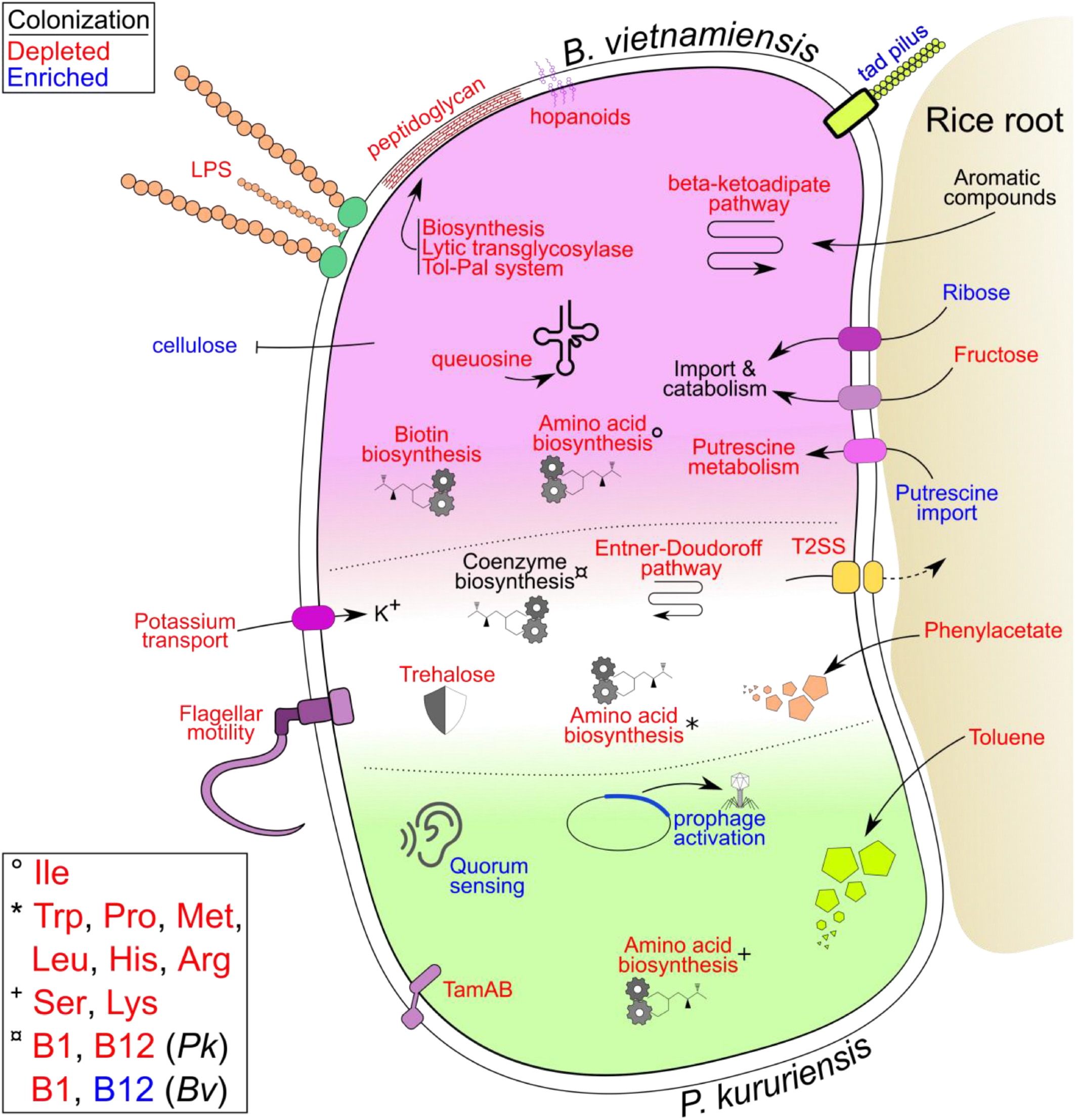
Summary diagram of colonization-impacted bacterial functions involved in efficient rice root colonization. Pathways, functions, and genes described throughout this work are synthetized on this integrative representation. Elements specific of *Bv* are placed in the top part of the schematic bacteria (pink) and *Pk* specific elements at the bottom (green). Elements that are commonly found in both bacteria are placed at the interface (white). Function and molecular systems which were detected to be colonization-depleted are written in red font whereas colonization-enriched genes are written in blue font.

### *Bv* and *Pk* have different strategies for rice root colonization

As was previously suggested by microscopy observations and host transcriptomics, *Bv* and *Pk* have different approaches to rice colonization (6). The present analysis highlights the respective genetic requirements of *Bv* and *Pk* during their rice colonizing process. Strikingly, a considerable higher number of genes is necessary for a successful colonization by *Bv* than by *Pk*, despite similar genome size. This requirement could result from increased plant defenses that *Bv* is exposed to compared to *Pk* (6). Supposedly, the pathogenic genetic background and potential of *Bv* could be responsible for this adverse plant response and necessitate a higher number of genes to act against. Moreover, as *Bv* is known to colonize multiple eukaryotic hosts, both plants and animals (41–44), we can also hypothesize that *Bv* expresses many genes which are not of use for plant colonization, thus a mutation in these genes will increase the bacteria’s fitness as the ensuing metabolic cost can be spared. This could explain that plant-detrimental genes were found in relatively similar amounts to plant-essential genes in *Bv* while they are considerably less frequent in *Pk* (**Figure 3A**). One striking example is the case of vitamin B12 production. This cofactor is among the most complex in bacteria and requires at least 19 enzymatic steps to produce while many enzymes further depend on B12 for their activity (45). Here, the synthesis of B12 is beneficial for the fitness of *Pk* during colonization while it has an adverse effect on *Bv*. This observation is supported both by the Tn-seq and the insertional mutagenesis approaches. We can hypothesize that the cofactor is involved in enzymatic activities that play a crucial role for colonization in *Pk* but not in *Bv*. As the production of B12 is associated with a high metabolic cost, sparing its expense could explain the fitness advantage displayed by *Bv* when the B12 synthesis pathway is mutated.

Defects in secretory activities are readily complemented in bacterial communities making the responsible secretion systems opaque for detection by Tn-seq approaches. Indeed, secretion systems are not detected by most studies relying on Tn-seq except when focusing on more sparsely colonized environment such as the apoplast (31). In the present study however, a T2SS is colonization-depleted in *Bv* and *Pk* and the type 6 secretion system (T6SS) in *Bv* during IR64 colonization. While the T6SS can be used for cell adhesion (46), a role beyond complementable secretion remains elusive for the T2SS. Still, as only one T2SS is supposedly involved in colonization it suggests a separate role from the second one present in *Pk*.

The only gene with an identified function that is predicted by the Tn-seq approach to be negatively involved in root colonization in both *Bv* and *Pk* is *pdeR,* Site-directed mutagenesis and a competition assay revealed that its mutation was highly beneficial for the root colonization activity of *Bv* as expected from the Tn-seq results, but detrimental for *Pk*, which was inconsistent with the Tn-seq analysis. PdeR is involved in the turnover of the secondary messenger c-di-GMP. In *Xanthomonas oryzae*, the deletion of *pdeR* results in a decreased virulence on rice (47). Through its association in two component signal transduction systems, PdeR could have a variety of indirect roles beside c-di-GMP cycling (48). The divergence observed between the Tn-seq and directed mutagenesis for *Pk* could also be linked to the inoculum’s concentration which is likely to have been superior in the latter approach and might have triggered a different plant response with the observed detrimental effects on the *Pk* mutant population (**Figure 6**).

### Tn-seq identifies common plant-colonization traits

To date, there is no transcriptome data available for *Bv* that would allow a comparison with our data. However, the transcriptomic response of *Pk* to rice root macerates has been assessed before (14). Out of the 471 colonization-depleted genes identified by the present approach, 267 are differentially expressed in *Pk* when stimulated with root macerate (**Table S5**). Dominantly, these genes are involved in amino acid metabolism, cell motility and cell wall/membrane biogenesis. The prevalence of these functions has further been reported by Tn-seq approaches in other root colonizing models such as *Dickeya dadantii* and *Pseudomonas spp.* (21, 23, 24). Our results are congruent with most plant-microbe colonization studies, underlining the importance of genes involved in the production of surface components for cellular attachment to the host (49). Our analysis also presents substantial overlap of colonization-depleted genes with what has been found in the plant growth promoting species *Azoarcus olearius* and *Herbaspirillum seropedicae* (50). Notably genes involved in peptidoglycan *(ampD)* and cell wall *(murAI)* formation, chemotaxis (*cheARW*), iron uptake (ferredoxins) and cobalamin synthesis (*cbiA*, *cobIO*) were detected.

Disruption of the ED pathway was demonstrated to reduce root colonization capacity in *Pseudomonas chlororaphis* with a subsequent loss of ISR stimulation (51). Later works on *Pseudomonas fluorescens* observed that the expression of *edd* was enhanced in the rhizosphere of *Arabidopsis* compared to liquid growing medium (52). Mutants for *edd* in this latter study also failed to stimulate ISR but without the colonization defect observed in *P. chlororaphis*. For pseudomonads, the importance of the ED pathway might be linked to its direct product, pyruvate, that is required for the synthesis of the ISR-eliciting butanediol (51). Other advantages might arise from the ED pathway such as the production of NAD(P)H which is not generated by the EMP glycolysis pathway. This cofactor is used by thioredoxins and could be involved in plant colonization and associated oxidative stress tolerance (33, 34).

### IR64 rice has more stringent requirements for colonization by *Bv* and *Pk* than Nipponbare rice

For both *Bv* and *Pk*, a stronger host-genotype effect on the number of colonization-impacted genes was observed during the association with IR64 compared to Nipponbare rice. For *Pk*, this genotype-effect is also observed on the root colonization profiles of the two cultivars (**Figure 2**). One specificity of *indica* rice (IR64) over *japonica* (Nipponbare) is its improved nitrate uptake and assimilation capacity (53), which is linked with the presence of the nitrate transporter NRT1.1B in the former, and proved to impact its microbiota (18). Nipponbare rice preferably imports the alternative nitrogen source ammonium, which can be correlated with the depletion of bacterial mutants impacted in the import of this nutrient (*amtB*) in the Nipponbare environment in our experimental set-up, as bacteria would compete with rice for the same nitrogen source) (**Table 3**).

It was interesting to observe that the T6SS, a major macromolecular system of *Bv*, was only colonization-depleted during inoculation on IR64 roots. The T6SS can be employed by bacteria for competition with other prokaryotes but also interaction with eukaryotes (54). As secretory function should be complemented by the bacterial community, the T6SS of *Bv* appears to be rather involved in adherence to eukaryotic cells, in a host-dependent manner.

Still, most colonization-impacted genes are conserved between rice cultivars and several functions such as motility, amino acid metabolism and biofilm production have been repeatedly described for their role in the general association of bacteria with plants (**Figure 7**). We have found evidence of an increased stress to which bacteria are exposed in the near vicinity of plants through the depletion of mutants involved in osmoprotection, toxic compound degradation and DNA reparation. The identification of several functions that are part of the core-genome shared by both bacteria but are only colonization-impacted in one strain reinforces the validity of these observations and the different adaptations that both bacteria must undergo during colonization of rice. This is especially true in the present system as *Bv* and *Pk* induce different levels of plant defenses (6).

An increasing amount of plant microbiome studies rely on Tn-seq to reveal the nature of genes underlying root colonization (20, 21, 23, 24, 31, 50). Tn-seq further offers the benefit over more common methods such as RNAseq, to inform on the genes obstructing colonization (mutants with higher colonization fitness), thus presenting a more complete catalogue of factors driving host-bacteria compatibility. Together with other trending methods such as microfluidic visualization technics and synthetic bacterial communities (49), Tn-seq proves here to be a powerful tool for a better understanding of the genetic bases underlying colonization of different hosts.

## Materials and Methods

### Bacteria and plant culture conditions

The strains used in this study are *Burkholderia vietnamiensis* LMG 10929 (*Bv*) and *Paraburkholderia kururiensis* M130 (*Pk*), either wild type, or modified by insertion of the pIN29 plasmid conferring chloramphenicol (Cm) resistance and DsRed fluorescence, or spontaneous rifampicin (Rif) and spectinomycin (Spt) resistant strains, or insertional mutants of a non-replicative plasmid in several candidate genes in each strain (**Table 4**) (6, 55). Spontaneous antibiotic resistant strains for LMG 10929 and M130 were obtained by plating 100 μL of bacterial liquid culture at OD_600_=1.0 on Luria’s low salt LB (LBm; Sigma-Aldrich) with either rifampicin (30 μg.mL^-1^) or spectinomycin (50 μg.mL^-1^). After 48h incubation at 28°C, single colonies were selected and streaked on fresh LBm plates with rifampicin or spectinomycin, then grown in broth LBm with antibiotics and stored in 20% glycerol at −80°C. From glycerol stocks, bacterial strains were cultured as follows: bacterial cells conserved at −80°C were plated on LBm agar plates (with antibiotic for mutants) and incubated for 72 h at 28°C. Single colonies were used to inoculate 10 mL of LBm broth (with antibiotic for mutants) in 50 mL Falcon tubes and incubated for various amounts of time allowing the different strains to reach an OD_600_=1.0. For inoculation purposes, cultures were adjusted to 5 x 10^7^ cfu.ml^-1^.

**Table 4.** Bacterial strains and plasmids used in this study.

| Strains/plasmids | Relevant characteristics and plasmid constructions | Reference/Source |
| --- | --- | --- |
| *E. coli* |  |  |
| MFDpir | Donor strain containing the pSAM-Ec vector and auxotroph for diaminopimelate, $\Delta dapA::(erm-pir)$ | (Ferrières et al., 2010) |
| *B. vietnamiensis* |  |  |
| LMG10929 | Wild-type strain | (Gillis et al., 1995) |
| Rif<sup>R</sup> | Spontaneous rifampicin resistance clone of LMG10929 | This study |
| Spt<sup>R</sup> | Spontaneous spectinomycin resistance clone of LMG10929 | This study |
| DsRed | LMG1029 + pIN29 | (King et al., 2019) |
| $\Omega cobW$ | Insertion mutant of pSHAFT2 in AK36_2246 | This study |
| $\Omega pdeR$ | Insertion mutant of pSHAFT2 in AK36_4666 | This study |
| $\Omega edd$ | Insertion mutant of pSHAFT2 in AK36_178 | This study |
| *P. kururiensis* |  |  |
| M130 | Wild-type strain | (Baldani et al., 1997) |
| Rif<sup>R</sup> | Spontaneous rifampicin resistance clone of M130 | This study |
| Spt<sup>R</sup> | Spontaneous spectinomycin resistance clone of M130 | This study |
| DsRed | M130 + pIN29 | (King et al., 2019) |
| $\Omega cobW$ | Insertion mutant of pSHAFT2 in ANSKv1_10390 | This study |
| $\Omega pdeR$ | Insertion mutant of pSHAFT2 in ANSKv1_51226 | This study |
| $\Omega edd$ | Insertion mutant of pSHAFT2 in ANSKv1_70910 | This study |
| Plasmids |  |  |
| pIN29 | Carries genes conferring DsRed fluorescence and chloramphenicol resistance. DsRed is under control of the highly active pTAC promoter | (Vergunst et al., 2010) |
| pSAM-Ec | Contains Himar1C9 transposase and a mariner transposon with a kanamycin resistance cassette | (Wiles et al., 2013) |
| pSHAFT2 | Vector used for insertional mutagenesis containing a chloramphenicol resistance gene (non-replicative plasmid) | (Shastri et al., 2017) |

*Oryza sativa subsp. japonica* cv. Nipponbare and *Oryza sativa subsp*. *indica* cv. IR64 were cultured as described in King et al. (2019). Briefly, seeds were sterilized using successive 70% ethanol and 3.6% sodium hypochlorite treatments and germinated seedlings were transferred onto sterile perlite in an airtight hydroponic system. 5-days old seedlings were inoculated with 1 mL of bacterial solution at 5 x 10^7^ cfu.ml^-1^ and grown up to 14 days at 28°C (16h light, 8h dark). For competition assays, five plants were inoculated with a mix at 1.10^7^ cfu.ml^-1^ containing 0.5 mL of SptR *Bv* or *Pk* strains (previously grown separately in broth LBm + Spt 50 μg.mL^-1^ and washed to remove antibiotic) and 0.5 mL of either one of the insertional mutants (Ω*cobW*, *ΩpdeR* or Ω*edd*) of the corresponding strain (grown in LBm + Cm 100 μg.mL^-1^ and washed to remove antibiotic).

### Estimation of root colonizing bacterial population

For the estimation of root colonizing populations, the plants were prepared as described above and inoculated with WT strains at 1 x 10^7^ cfu.ml^-1^. Three systems (15 plants) were prepared for each condition and grown for 1, 7 or 14 days (28°C; 16 h light; 8 h dark). At each time point, five plants were harvested for each condition. The entire root systems were sampled and separately transferred to screw cap tubes containing 1 mL of sterile water and a sterile ceramic bead. Roots were weighted and then pulveri1zed using a Fastprep-24 (MPbio) at 6 m.s^-1^ for 40 seconds. A serial dilution of the resulting solution was spotted out on square LBm plates and incubated at 28°C for 48 hours. The number of colony forming units (cfu) were counted and adjusted to the weight of the root systems. The mean colonization values were compared between cultivars and between bacteria genotypes at each time point and for each bacterium separately using a Wilcoxon test. Results were considered significantly different at p<0.05. For the competition assay between WT and mutant strains, plants were harvested at 7 dpi and processed as described above, except the bacterial dilutions were spotted on LBm plates containing either streptomycin (30 μg.ml^-1^) or Cm (200 μg.ml^-1^) to select respectively the WT or the mutant strains.

### Tn-seq library preparation

A Himar1 mariner transposon carried by the pSAM_EC vector in *Escherichia coli* strain MFDpir (56, 57)was introduced into *Paraburkholderia kururiensis* M130 Rif^R^ and *Burkholderia vietnamiensis* LMG 10929 Rif^R^ through conjugation. Both donor and recipient were grown until OD_600_=1.0 in liquid low-salt LB (LBm) and the medium of MFDpir was further supplemented with diaminopimelic acid (DAP; 300 μg.mL^-1^). Cells were spun down and washed once with LBm and concentrated to a final OD_600_=50. Donor and recipient strains were mixed 1:1 and 50 μL suspensions were spotted on square Petri dishes containing LBm and DAP. Growth rates were different for both recipient species and conjugation time was adapted accordingly. The mating mix was incubated for 6 h and 24 h at 28°C for LMG 10929 and M130, respectively. The spots where then resuspended in 2 mL LBm per plate. The mating mix was further spread on LBm Petri dishes with rifampicin (30 μg.mL^-1^) and kanamycin (50 μg.mL^-1^) and incubated at 28°C. This positively selects for recipient strains having integrated the transposon and negatively selects for the DAP auxotroph *E. coli* donor. After growth, the colonies were resuspended in 1 mL LBm per plate. The library was separated into 1 mL aliquots, and stored in 20% glycerol at −80°C. Additionally, a serial dilution of the mating mix followed by spreading on LBm with rifampicin (30 μg.mL^-1^) and kanamycin (50 μg.mL^-1^) was used to estimate the abundance of mutants in the library through cfu counting.

### Tn-seq experimental setup

Tn-seq libraries were thawed on ice and diluted in sterile water to OD_600_=1 and then inoculated in 50 mL liquid LBm with rifampicin (30 μg.mL^-1^) and kanamycin (50 μg.mL^-1^) at a final OD_600_=0.1 and grown at 28°C 150 rpm until OD_600_=1.0. One part was conserved for plant inoculation while the other was centrifuged at top speed for 10 minutes and flash frozen for subsequent DNA extraction to serve as control condition. Plants were grown in hydroponic systems as described above. After 5 days of growth, each plant was inoculated with 1 mL Tn-seq library bacterial suspension at 5 x 10^7^ cfu.mL^-1^. The experimentation was performed in triplicates with each replicate consisting of five plants. At 7 dpi, the roots were harvested and placed in TE buffer.

### DNA extraction and sequencing methods

The bacterial genomic DNA isolation using CTAB protocol (JGI) was used for the extraction of bacterial DNA from the control and experimental conditions. For the latter, whole roots were used in the first stages of the protocol. Whole roots were immersed in the extraction solution and vortexed for 5 min at each step of the protocol to facilitate bacterial separation. Root grinding was avoided to prevent excessive DNA contamination from the plant material. Roots were removed from the extraction buffer after the lysis steps, before CTAB is added.

Ten μg of total DNA were digested with 1 μL of MmeI restriction enzyme (New England Biolabs) in 250 μL total volume with 10 μL of S-adenosyl methionine (SAM) 1.5 mM and 25 μL CutSmart buffer during 1 h at 37°C. Then, 1 μL of FastAP Thermosensitive Alkaline Phosphatase 1 U.μL^-1^ (ThermoScientific) was added and incubated 1h at 37°C. All enzymes were inactivated through a 5 min incubation at 75°C. Digested DNA was column purified using a NucleoSpin Gel and PCR Clean-up kit (Macherey-Nagel). 2 μg of purified DNA was ligated to adapters containing specific barcode sequences. Adaptors were obtained by annealing the primers 5’-TTCCCTACACGACGCTCTTCCGATCTXXXXXNN-3’ and 5’-YYYYYAGATCGGAAGAGCGTCGTGTAGGGAAAGAGT-3’ where NN are random nucleotides for annealing to the dinucleotide overhang generated by MmeI and XXXXX and YYYYY are complementary, barcode-specific sequences. Ligations were performed using 1 μL of 5 mM adapters, 1.5 μL of T4 DNA ligase 1 Weiss U.μL^-1^ (ThermoScientific) and 2.5 μL 10x ligase buffer in a total volume of 25 μL and incubated overnight at 16°C. 1 μL of ligation product was amplified using PCR with a GO Taq DNA polymerase (Promega) and Illumina primers (P7 5’-CAAGCAGAAGACGGCATACGAGATAGACCGGGGACTTATCATCCAACCTGT-3’; P5 5’-AATGATACGGCGACCACCGAGATCTACACTCTTTCCCTACACGACGCTCTTCCGATCT-3’). 22 PCR cycles were run (30 s at 92°C; 30 s at 60°C; 1 min at 72°C) with an initial step at 92°C for 2 min and a final step at 72°C for 10 min. PCR products were subjected to a final gel purification using the NucleoSpin Gel and PCR Clean-up kit (Macherey-Nagel).

Sequencing was performed at the high-throughput sequencing platform at I2BC (CNRS, Gif-sur-Yvette, France) using Nextseq500 (Illumina) technology and 80 sequencing cycles (single read). Preliminary data analysis (demultiplexing, trimming and mapping) was performed by the sequencing platform. Sequences were trimmed from barcodes and mariner transposon sequences and deposited in European Nucleotide Archive under Bioproject number PRJEB42565.

### Tn-seq data analysis

Gene essentiality for optimal growth in a liquid LBm broth under agitation was assessed through a hidden Markov model (HMM)-based analysis using the EL-ARTIST pipeline (window size of 7 TA sites; *P* value of 0.03) (58). The pipeline predicts whether a TA site is located in an essential or nonessential region. In some cases, the locus is predicted to be located in a region that contains both domains that are required and dispensable for growth. After the first refinement, the HMM will recalculate the transition probabilities based on the new data and repeat the run until the algorithm reaches convergence.

For the identification of conditionally essential genetic regions, the sorted aligned sequences were first normalized using TRANSIT 3.1.0 (59) with the Trimmed Total Reads method at default settings and an additional LOESS correction. The normalized datasets were then analyzed using the TnseqDiff function of the R package Tnseq (60). Adjusted p-values were calculated using the Benjamini & Hochberg correction. Genetic loci were considered colonization depleted when their occurrence in the experimental condition was at least 1.5-fold lower than in the control condition at a confidence level of padj<0.05 and inversely for colonization enriched genes.

### Comparative genomics procedures

Core genome compositions were calculated using the Phyloprofile exploration tool implemented in the MicroScope microbial genome annotation and analysis platform (61). Homology constraints were set at minLrap ≥ 0.8 and identity ≥ 50%. We used the COGNiTOR pre-computed COG category classification available on MicroScope to infer COG category affiliations of *Bv* and *Pk* genes.

### Construction of insertional mutants

For the production of the *cobW, pdeR* or *edd* insertional mutants of *B. vietnamiensis* LMG10929 and *P. kururiensis* M130, a gene fragment of either *cobW, pdeR* and *edd* of each strain was amplified using specific primers (**Table S6**). Genes fragments were inserted into the target vector pSHAFT2 (Shastri et al., 2017) carrying a chloramphenicol (Cm) resistance gene using an XbaI/XhoI double digestion. Insert and plasmid were mixed at a 5:1 ratio with regard to their respective molecular size, ligated using a T4 DNA ligase (Promega) and cloned into heat-shock competent *E. coli* DH5α cells. Positive cells were multiplied and their plasmid extracted using the Wizard Minipreps DNA purification system (Promega). Plasmids were electroporated into WT *B. vietnamiensis* LMG10929 and *P. kururiensis* M130 strains. Transformed cells were transferred in LBm for 3h at 30°C and then spotted on LBm plates containing Cm (200 μg.ml^-1^) for selection of mutants. Plasmid insertion in each targeted gene was checked by PCR. Mutants and WT strains showed identical growth rates in broth LBm or Hoagland medium supplemented with glucose (not shown).

## Acknowledgments

The authors acknowledge the IRD itrop HPC (South Green Platform; https://bioinfo.ird.fr – http://www.southgreen.fr) at IRD Montpellier for providing HPC resources that have contributed to the research results reported here. We acknowledge the High-throughput sequencing facility of I2BC for its sequencing and bioinformatics expertise. The LABGeM (CEA/Genoscope & CNRS UMR8030), the France Génomique and French Bioinformatics Institute national infrastructures are acknowledged for support within the MicroScope annotation platform.

## Funding

The authors gratefully acknowledge financial support from the CGIAR research program (CRP) RICE as well as from the French national research agency (ANR) funding the BURKADAPT project (ANR-19-CE20-0007). AW, JL and LG were supported by PhD fellowships from the French Ministry of Higher Education, Research and Innovation.

